# GENOMIC PROFILING OF SARS-COV-2 STRAINS CIRCULATING IN SOUTH EASTERN REGION OF INDIA DURING THREE WAVES OF PANDEMIC

**DOI:** 10.1101/2024.01.06.574128

**Authors:** Vanathy Kandhasamy, Agieshkumar Balakrishna Pillai, Vignesh Mariappan, Malarvizhi Ramalingam, Pajanivel Raganadin, Ramya Ramadoss, Balasubramanian Moovarkumudalvan, Joshy M Easow, Madavan Vasudevan, Rao S.R.

## Abstract

Continuous bio-surveillance of SARS-CoV-2 is an ongoing task at local, national and global levels since the pandemic onset for understanding genetic evolution and vaccine efficacy. Present study was designed to track the emergence of new variants along the duration of three peaks of infection in the city of Puducherry, India. A total of 128 samples were subjected to Illumina deep RNA sequencing. The results showed predominance of uncommon, delta and omicron variants in first, second and third waves respectively. The most common pangolin lineage was B.1.560 and B.1.617.2. The study observed a total of 3133 common and 11 new mutations. The most common is in the Spike_D614G. A new set of mutations was observed in key viral factors such as NS16 that are implicated to be involved in immune evasion. This may have impact on enhanced disease virulence, vaccine efficiency and possible tolerance to current antivirals. This warrants further in vitro studies to understand the significance of the mutations. While the results presented would also augment the ongoing research on evolutionary and the genetic epidemiology of SARS-CoV-2, it also emphasizes the need for continuous genetic monitoring to predict the forthcoming threats due to the emergence of new or existing variants.

## Introduction

The Coronavirus disease-19 (COVID-19) caused by severe acute respiratory syndrome-coronavirus-2 (SARS-COV-2) first originated in Wuhan city of China by the end of 2019. World Health Organization (WHO) declared COVID-19 as pandemic on March 11^th^, 2020 (Cucinotta & Vanelli, 2020). In India, the first case was reported in January 2020. Since then, three waves of pandemic were reported with different variants ranging from mild to severe infection (Mandal et al., 2021). The current number of cases had reached 4.44 crores and 5.3 lakhs deaths in India as per Ministry of Health and Family Welfare as on 13^th^ July 2023. In the Union territory (UT) of Puducherry, the first occurrence of cases was seen on 17^th^ March 2020 and currently reached up to 0.166 million cases and 1962 deaths all over the UT. The virus had mutated over a period into different variants which confer increase in severity of infection, rapid transmission and immune escape which is of more concern (Harvey et al., 2021). Bio surveillance involves the early detection of pathogens and variants for mitigation efforts and limit its spread through continuous genome sequencing approaches (Bartlow et al., 2021). The genomic surveillance revealed that there are multiple lineages circulating in India and in other countries (Sarkar et al., 2021). The ancestral lineages of Phylogenetic Assignment of Named Global Outbreak (PANGO) classification system are A and B. The variants initially responsible for outbreak in Italy belong to B.1 and it had an amino acid change in spike protein D614G. Subsequently there had been several mutation in B.1 with replacement of one or two amino acid resulting in emergence of several other variants of concern (VOC) such as B.1.1.7 (N501Y-Alpha), B.1.351 (N501Y,E484K,K417T- Beta), P.1 from the lineage B.1.1.28 (N501Y,E484K,K417T-Gamma), Delta variant (B.1.617.2) from United Kingdom, South Africa, Brazil and India respectively (Faria et al., 2021; Frampton et al., 2021; Tegally et al., 2021). These variants had several implications on public health regarding the severity, hospitalisation, mortality and immune escape to vaccination and therapeutic antibodies (Harvey et al., 2021). The genomic surveillance helps in early identification of the emerging VOC and in implementation of public health measures. There are also several variants of interest (VOI) circulating such as Epsilon (B.1.427), Theta (P.3), Iota (B.1.526), Kappa (B.1.617.1) (Thakur et al., 2022). The current dominating strain circulating in many countries is Omicron variant (B.1.1.529) which was reported from South Africa on November 2021. Because of its high transmissibility, immune escape, and high rate of transmission among vaccinated individuals, it has spread rapidly across the globe (Khandia et al., 2022). WHO declared Omicron sub lineage BA.2 as the dominant strain. There are few amino acid differences between BA.1 and BA.2. The studies found that BA.2 is more contagious and increase in reinfection rate compared to BA.1 (Lyngse et al., 2022; Sahebi & Keikha, 2022). So, there is a close monitor on this lineage to know more about its severity and pathogenesis. Continuous bio-surveillance and analysis of sequences from cases from various affected parts of the country would provide knowledge on genetic evolution and mutation rate of the virus. In India, the first two SARS-CoV-2 sequences were found to have two different introductions into the country. Later delta type was first identified in Maharashtra which outcompeted the previously existing strains like alpha and beta (Yadav et al., 2020). Having vaccines in usage in National levels, rapidly evolving mutation rates and any minor modifications in the viral genome may make vaccines more or less effective emphasis the need to monitor the viral lineages and the variants in circulation for effective surveillance. In this context, the present study was designed to track the variants, emergence of new lineages between three peaks of infection in Puducherry, a coastal region in South Eastern India. The study was carried out using the RNA sample obtained from the swab samples collected from the National Accreditation Board for testing and Calibration Laboratories (NABL) accredited COVID-testing facility in a tertiary care hospital (Coumare et al., 2021) and the results obtained based on novel and known mutations are discussed in the study.

## Methods

### Study approval

The study was approved by the Institute Research Committee and Institute Human Ethics Committee - MGMCRI/IRC/52/2021/04/IHEC/42. All the methods were performed in accordance to the Helsinki guidelines.

### Nucleic acid extraction, COVID-19 screening and sample selection

Oropharyngeal/Nasopharyngeal samples received in viral transport medium at the diagnostic facility for molecular detection of RNA virus were used for the study. The host institute has started its Real-Time Reverse Transcriptase PCR (RT-PCR) testing service in August 2020 after obtaining NABL accreditation as per Indian Council of Medical Research (ICMR) recommendation for COVID testing. A total of 10,561 samples were processed from August 2020 to February 2022. Among them 1699 were positive. The whole genome sequencing (WGS) was done in representative positive samples (n=128).

RNA was extracted from the nasopharyngeal/oropharyngeal samples using QIAamp RNA extraction kit and stored at -80°C. RT-PCR was done using various ICMR approved COVID-19 RT-PCR kits. For WGS, samples that are positive for SARS-CoV-2 by RT-PCR along with the CT value ≥18 & ≤25; State of Residence – Puducherry were used as the criteria for sample selection. A total of 128 samples fulfilling the above criteria were selected which included 17 from first wave (September 2020 to November 2020), 80 from the second wave (March 2021 to August 2021) and 31 from third wave (September 2021 to February 2022).

### Retrieval of demographic and clinical details of patients

The demographic details were collected from ICMR specimen reference form. Since most of the forms were partially filled due to case load; the necessary details such as vaccination status, breakthrough infection (individuals turning positive for COVID-19 infection after 14 days of vaccination), co-morbidities, home quarantined or hospitalised during the course of infection were collected with an informed consent.

### Library Preparation

RNA samples where in CT values ≥18 & ≤25 was considered for the WGS. The RNA was converted into in cDNA using a balanced panel sets of random hexamers, Oligo DT’s and COVID-19 specific oligos. SARS-CoV-2 genome is selectively amplified with oligo sets custom designed based on the ARTIC protocols. These samples are purified and barcode is added through a barcoding PCR cycle. The final samples are purified, quality checked, quantified and equimolar pooled for sequencing using Illumina Miseq deep sequencing platform with 300 X 2 bp Paired End chemistry. 1000x of the genome size raw data was generated to ensure maximum coverage and depth for each sample. Illumina MiSEQ platform with 300x2 was preferred for COVID-19 genome sequencing for reasons of throughput, base quality and the supported bioinformatics pipelines.

### Deep sequencing data quality control and consensus genome assembly

Raw data quality control followed by adapter trimming was done using FASTQC tool kit, Trim Galore and Cut adapt. The processed reads are used to align to the reference strain NC_045512.2 using BWA and SamTools/Pileup to calculate coverage and depth. The consensus sequences are derived from the alignment files. This consensus genome sequence was used as input for identification and classification of Pangolin and Nextclade lineages.

### Identification and Characterization of SARS-COV-2 isolates

Consensus draft genomes of COVID-19 isolates were subjected to various analysis using CoVsurver (https://gisaid.org/database-features/covsurver-mutations-app/), NextClade (https://clades.nextstrain.org/), Vigor (https://www.viprbrc.org/brc/vigorAnnotator.spg) and Genome Detective (https://www.genomedetective.com/app/typingtool/cov/). The results from the analysis tools were analysed using Microsoft Excel and GraphPad Prism tool for visualization purposes. The resulting consensus genomes (OP599771-OP599898) have been deposited to GenBank and the raw reads have been submitted to the NCBI-Sequence Read Archives (SRA) with submission ID (SUB12132093). The phylogenetic relationship between sample consensus genome sequences was analyzed using IQ-Tree tool (version 2.2.2.3) (Minh et al., 2020) by constructing a phylogenetic tree using 1000 bootstrap iterations and was visualized and annotated using iTOL online tool (Letunic & Bork, 2021)

## Results

### Demographic Profile of the study participants

A total of 128 samples (17 samples from first wave, 80 samples from second wave and 31 samples from third wave) were sequenced which belongs to the age group of 20 to 87 years with an average age of 46.3 years. The age distribution was 20-30 years - 22 cases, 31- 40 years - 35 cases, 41-50 years - 23 cases and ≥51 years - 48 cases. Among these cases, the number of males were 79/128 (61.7%) and females 49/128 (38.2%). The break through infection after vaccination was found in 8 patients (among them three were hospitalised in which two patients had associated Diabetes mellitus). The comorbidity like diabetes mellitus/hypertension/hyperlipidaemia/heart disease were seen in 17 patients. Those patients under home quarantine had no comorbidity except 3 patients who had diabetes mellitus/hypertension. The fatality was around 4 cases in first wave, 5 cases in second wave and no cases in third wave. Their age group varied from 45-82 years. Around 81% of cases were individuals who wish to get themselves tested. The remaining are those with high risk, severe acute respiratory infection (SARI), those with Influenza like illness and asymptomatic contacts. Clinical and demographic details of the study subjects are given in **Table 1**.

**Table 1.**
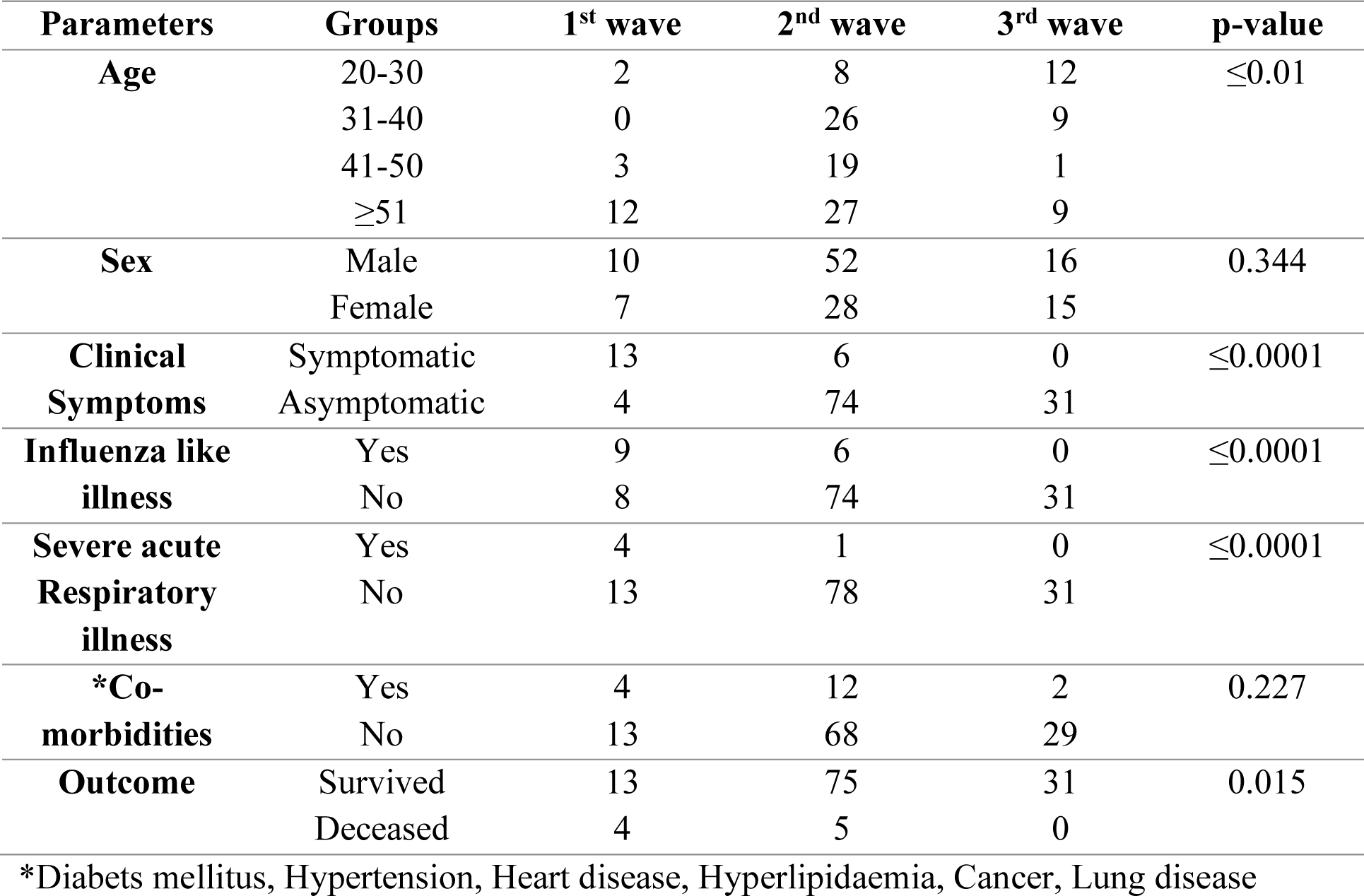
Clinical and demographic details in all three waves.

| Parameters | Groups | 1<sup>st</sup> wave | 2<sup>nd</sup> wave | 3<sup>rd</sup> wave | p-value |
| --- | --- | --- | --- | --- | --- |
| **Age** | 20-30 | 2 | 8 | 12 | $\leq 0.01$ |
|  | 31-40 | 0 | 26 | 9 |  |
|  | 41-50 | 3 | 19 | 1 |  |
| | $\geq 51$ | 12 | 27 | 9 | |
| **Sex** | Male | 10 | 52 | 16 | 0.344 |
|  | Female | 7 | 28 | 15 |  |
| **Clinical Symptoms** | Symptomatic | 13 | 6 | 0 | $\leq 0.0001$ |
|  | Asymptomatic | 4 | 74 | 31 |  |
| **Influenza like illness** | Yes | 9 | 6 | 0 | $\leq 0.0001$ |
|  | No | 8 | 74 | 31 |  |
| **Severe acute Respiratory illness** | Yes | 4 | 1 | 0 | $\leq 0.0001$ |
|  | No | 13 | 78 | 31 |  |
| **\*Co-morbidities** | Yes | 4 | 12 | 2 | 0.227 |
|  | No | 13 | 68 | 29 |  |
| **Outcome** | Survived | 13 | 75 | 31 | 0.015 |
|  | Deceased | 4 | 5 | 0 |  |

\*Diabetes mellitus, Hypertension, Heart disease, Hyperlipidaemia, Cancer, Lung disease

Sequencing results identified SARS-CoV-2 strains that could be classified under VOC, VOI and uncommon category. Out of 128 samples more than half turned out to be VOC, whereas another major slice was identified as uncommon strains as per WHO classification. The below pie chart shows the percentage/distribution of variants under each category based on the sequencing results **(Fig. S1)** and the total number of cases by variants are shown in **Table S1**. Genome coverage analysis of all the sequenced strain indicates uniform coverage of all the strains as illustrated in the **Fig. S2**.

### Distribution of SARS-COV-2 lineages

A complete description of all the lineages and the variants sequenced are provided in **Table 2** and the individual lineage distribution is given in **Fig. 1**. The most common among them is B.1.617.2 (n=64, 50%) followed by BA.2 (n=15, 12%) and B.1.560 (n=14, 11%). But there are also other circulating lineages which waned over time. During the first wave the most predominant lineage seen in our sequenced sample is B.1.560 (n=14), in second wave 6 different lineages were identified, of which the most common one being B.1.617.2 (n=61) followed by B.1.1.7 (n=5) and in third peak it is BA.2 (n=15) outnumbered the other lineages and 3 of the samples had overlapping B.1.617.2 of second wave. Among the symptomatic ILI and SARI individuals (N=12/19, 63.1%), majority were B.1.560 lineage. The common lineages among deceased individuals were B.1.560 (n=3), B.1.617.2 (n=4) and B.1.1.7 (n=2).

**Fig 1.**
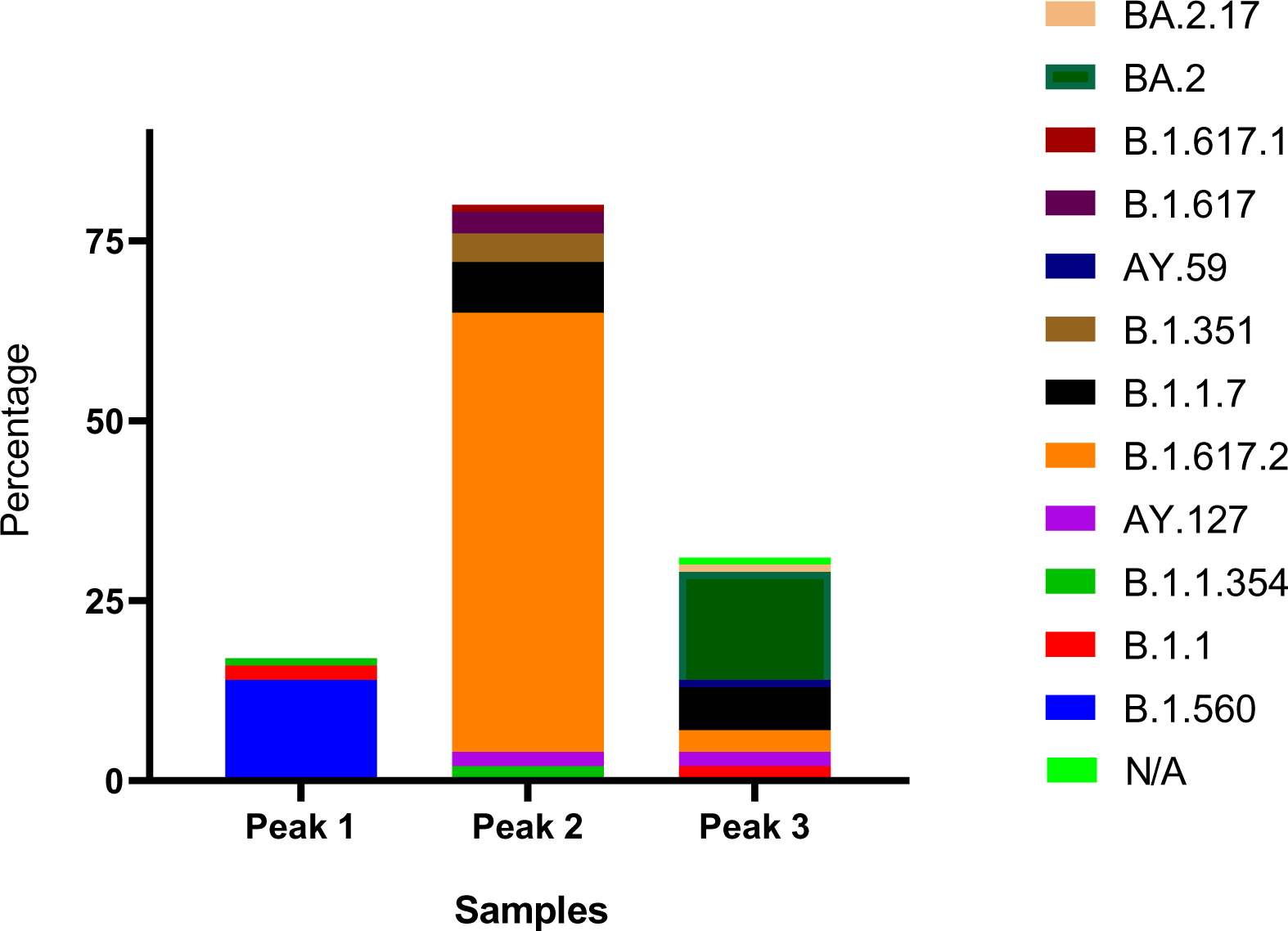
Individual lineage distribution during three major peaks

**Table 2.** Genome Classification.

| Lineage | Number of positives | Description | Wave | Variant |
| --- | --- | --- | --- | --- |
| **B.1.560** | 13 | Tokyo, Japan on 20200109 | First | UNCOMMON |
| **B.1.560** | 1 | Victoria, Australia on 20200230 | First | UNCOMMON |
| **B.1.1** | 2 | Tokyo, Japan 20200109 | First | UNCOMMON |
| **B.1.1.354** | 1 | Tokyo, Japan 20200109 | First | UNCOMMON |
| **AY.127** | 1 | North Rhine-Westphalia, Germany on 20200511 | Second | UNCOMMON |
| **AY.127** | 1 | Jammu and Kashmir, India on 20210321 | Second | UNCOMMON |
| **B.1.1.354** | 2 | Tokyo, Japan 20200109 | Second | UNCOMMON |
| **B.1.1.7** | 2 | California, USA on 20200708 | Second | VOC_Alpha |
| **B.1.1.7** | 5 | North Dakota, USA on 20201023 | Second | VOC_Alpha |
| **B.1.351** | 4 | Gauteng, South Africa on 20200801 | Second | VOC_Beta |
| **B.1.617.2** | 2 | California, USA on 20210106 | Second | VOC_Delta |
| **B.1.617.2** | 1 | Dakar, Senegal on 20200512 | Second | VOC_Delta |
| **B.1.617.2** | 1 | England, United Kingdom on 20210406 | Second | VOC_Delta |
| **B.1.617.2** | 2 | Gujarat, India on 20210407 | Second | VOC_Delta |
| **B.1.617.2** | 1 | Indiana, USA on 20210418 | Second | VOC_Delta |
| **B.1.617.2** | 23 | Jammu and Kashmir, India on 20210321 | Second | VOC_Delta |
| **B.1.617.2** | 12 | North Rhine-Westphalia, Germany on 20200511 | Second | VOC_Delta |
| **B.1.617.2** | 2 | Oromia, Ethiopia on 20210331 | Second | VOC_Delta |
| **B.1.617.2** | 4 | Podravska, Slovenia on 20201204 | Second | VOC_Delta |
| **B.1.617.2** | 1 | Sicily, Italy on 20210316 | Second | VOC_Delta |
| **B.1.617.2** | 2 | Texas, USA on 20200716 | Second | VOC_Delta |
| **B.1.617.2** | 2 | Not Associated | Second | VOC_Delta |
| **B.1.617.2** | 1 | Diourbel, Senegal on 20200327 | Second | VOC_Delta |
| **B.1.617.2** | 1 | Gujarat, India on 20210401 | Second | VOC_Delta |
| **B.1.617.2** | 2 | Minnesota, USA on 20200527 | Second | VOC_Delta |
| **B.1.617.2** | 1 | Minnesota, USA on 20201009 | Second | VOC_Delta |
| **B.1.617.2** | 1 | Mizoram, India on 20210213 | Second | VOC_Delta |
| **B.1.617.2** | 2 | Red Sea Governorate, Egypt on 20201213 | Second | VOC_Delta |
| **B.1.617** | 2 | Delhi, India on 20200303 | Second | VOI_Kappa |
| **B.1.617** | 1 | Jharkhand, India on 20210106 | Second | VOI_Kappa |
| **B.1.617.1** | 1 | Jharkhand, India on 20210106 | Second | VOI_Kappa |
| **AY.59** | 1 | Not Associated | Third | VOC_Delta |
| **B.1.617.2** | 2 | Mecklenburg-Western Pomerania, Germany on 20210503 | Third | VOC_Delta |
| **B.1.617.2** | 1 | Not Associated | Third | VOC_Delta |
| **AY.127** | 1 | North Rhine-Westphalia, Germany on 20200511 | Third | VOC_Delta |
| **AY.127** | 1 | Not Associated | Third | VOC_Delta |
| **BA.1.1** | 2 | Not Associated | Third | Omicron |
| **BA.1.1.7** | 4 | Not Associated | Third | Omicron |
| **BA.1.1.7** | 1 | Lambayeque, Peru on 20220117 | Third | Omicron |
| **BA.1.1.7** | 1 | Tennessee, USA on 20220120 | Third | Omicron |
| **BA.2** | 1 | Karnataka, India on 20220117 | Third | Omicron |
| **BA.2** | 10 | Karnataka, India on 20220119 | Third | Omicron |
| **BA.2** | 1 | Santa Fe, Argentina on 20220405 | Third | Omicron |
| **BA.2** | 3 | Not Associated | Third | Omicron |
| **BA.2.17** | 1 | Karnataka, India on 20220117 | Third | Omicron |
| **#N/A** | 1 | California, USA on 20211215 | Third | Omicron |

### Breakthrough Infections

Based on the telephonic conversation the status of vaccinated individuals was recorded. Out of 128 samples, 105 responded to their vaccination status. Of which, only 6% were found to be vaccinated at the time of sampling. Most of the vaccinated individuals were found to harbour uncommon variants (n=3) followed by alpha, beta, kappa and omicron (n=1 each) as shown in **Fig. S3**. An overview of the variants sequenced along with the outcome is shown in **Fig. S4**.

### Detection of lineages and their evolutionary relationship

A total of 14 Lineages were identified by PANGO lineage. According to Nextclade assignment, the phylogenetic relationship of all detected lineages falls under 10 different clades. Similarly, CoVsuver and WHO analysis showed 7 different clades. The classifications of lineages based on pangolin, Nextclade, CoVsuver and WHO analysis are shown in **Fig 2** **A-D**.

**Fig 2.**
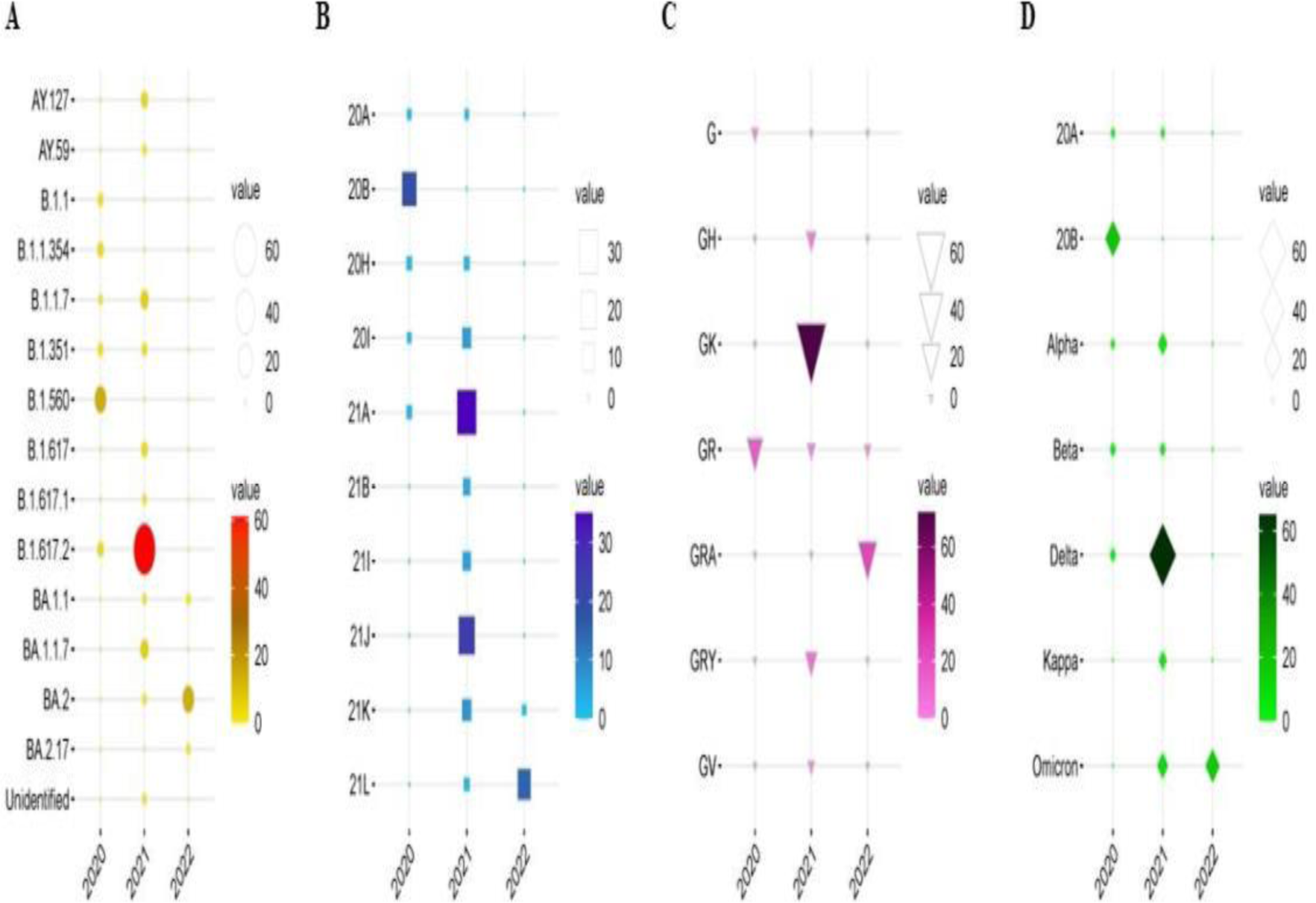
Classification of lineages and number of strains sequenced **(A)** Classification of lineages based on Pangolin; **(B)** Classification of lineages based on Nextclade; **(C)** Classification of lineages based on CoVsuver; **(D)** Classification of lineages based on WHO. The X-axis indicates the pandemic wave by year and Y axis indicates various clades/lineage/strain/type. The size of the objects viz circle, square, triangle and diamond indicate the total number of patients infected.

The study used FastTree-2 maximum-likelihood phylogeny to determine the phylogenetic relationships between the samples. FastTree-2 uses maximum-likelihood nearest-neighbor interchanges (MLIs), minimum-evolution subtree-pruning-regrafting (SPRs), and nearest-neighbor interchanges (NNIs). Our samples were aligned against the Wuhan reference genome (NC_045512.2). The branches of pangolin, Nextstrain and WHO lineages are depicted in **Fig. S5 A-C.**

### Mutational divisions of SARS-COV-2 detected during three peaks

We studied the frequency of mutations with known and new set of mutations. The total number of mutations 3144 (3133 common mutations and 11 new mutations) with a median of 24 mutations per sample was noticed. The greatest number of mutations were found in S gene (32% of total mutations) followed by N (12%) and NSP3 (7%) gene. The distribution of mutations in the sequenced strains are given in **Fig. 3**. The most common mutations observed was Spike_D614G. It is present across all the three peaks. The number of mutations per sample during first peak was found to be 8 whereas it increased gradually in further peaks resulting in 23 per sample in second and 40 per sample in the third peak. The phylogenetic relationship between sample consensus genome sequences could be inferred from phylogenetic tree (**Fig. 4**), which also provides an overall summary of sample attributes, predicted pangolin lineage, clade classification by CoVsuver and the unique mutations detected: NS7a_I4J, NSP16_I171T, NSP3_I1094S, NSP3_S925Y, NSP4_T370A,

**Fig 3.**
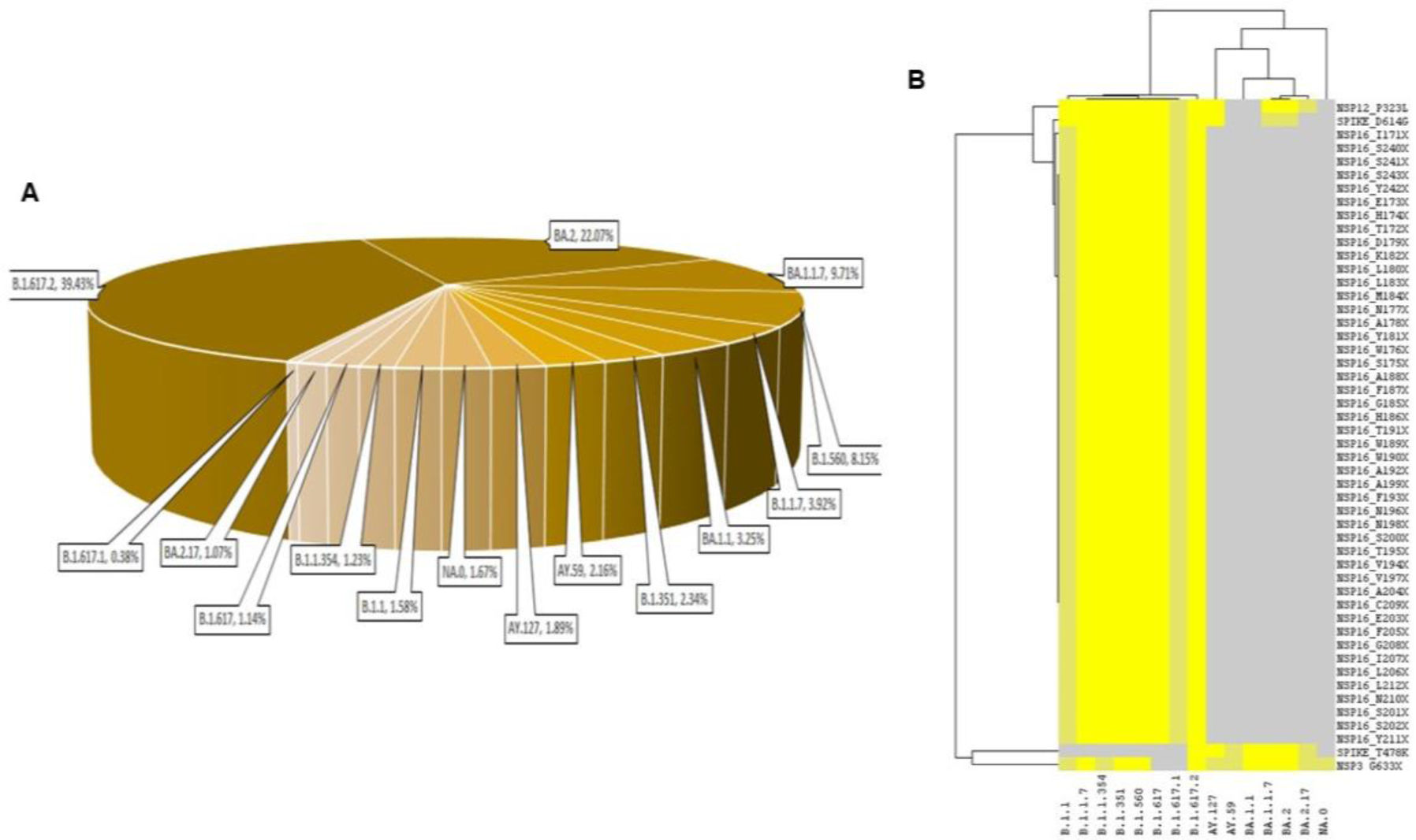
Distribution of Mutations **(A)**Mutation percentage in the sequenced lineages; **(B)** Dendrogram revealing the association between different mutation and sequenced strains

**Fig. 4.**
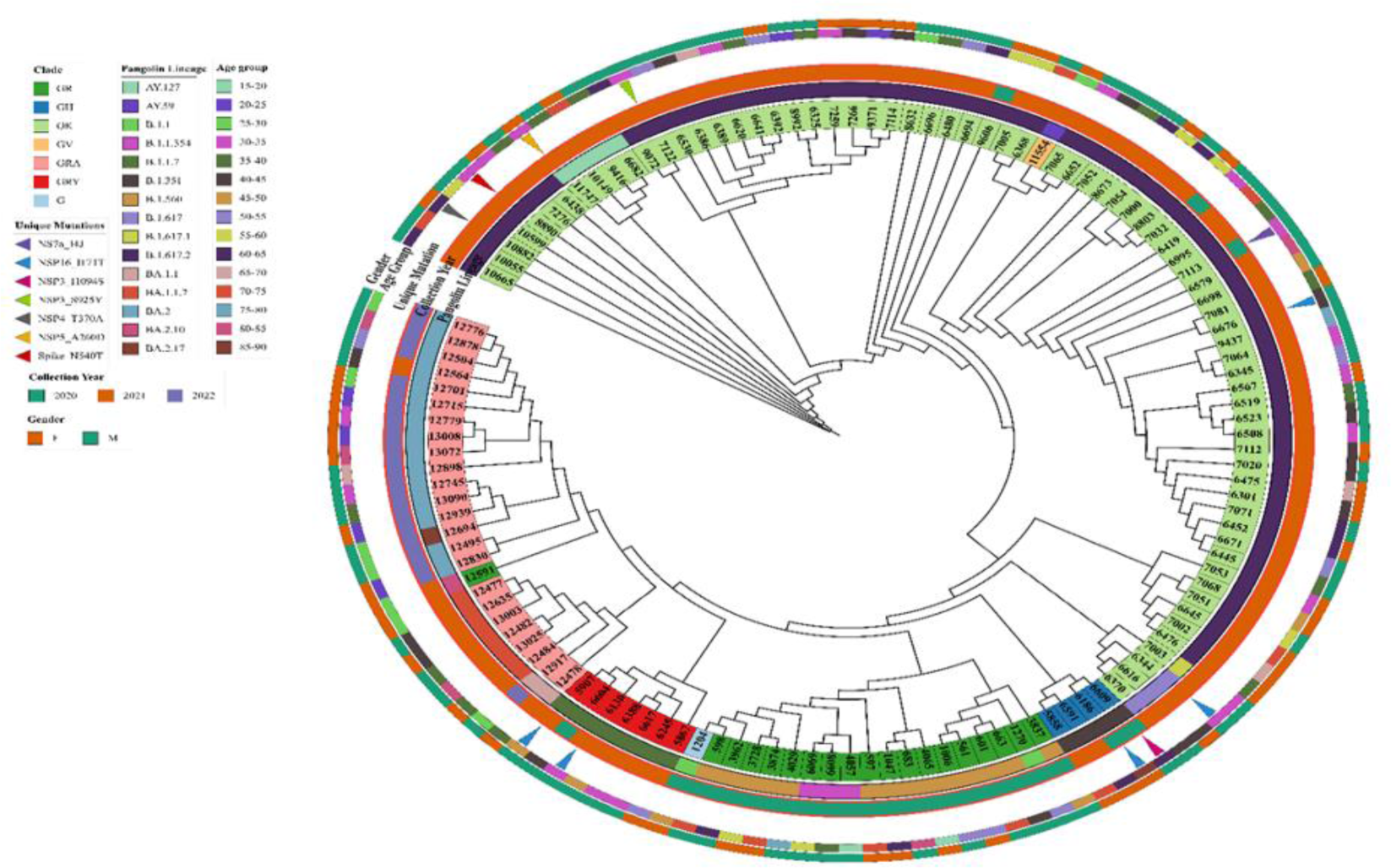
Phylogenetic tree of the consensus genome sequences along with sample attributes Tree labels are highlighted based on clade classification by CoVsuver as described in the legend. Lineages based on Pangolin are represented by the color strip 1. The corresponding sample collection year is represented by color strip 2. Samples with unique mutations are indicated by colored triangles. The patient age group and gender details are represented by color strips 3 & 4 respectively.

NSP5_A260D and Spike_N540T. Most of the unique mutations were detected from samples collected during second wave. The unique mutation NSP16_I171T was detected among 5 samples. Which is the greatest set of samples to contain a unique mutation. Except for one sample, the majority of samples were from male patients in the age group 30-50. Moreover, the detected unique mutation is quite significant since the mutation bearing consensus genome sequences were mostly estimated to have good coverage (>90%) and quality control score. The overall distribution of mutants aligning with the three major peaks as reported by Government of Puducherry (https://covid19dashboard.py.gov.in/) is represented in **Fig S6.**

## Discussion

The WHO declared COVID-19 as a pandemic disease on March 11, 2020. The first case was reported from India on January 30, 2020 of a patient with a travel history from Wuhan, China (Cucinotta & Vanelli, 2020). Since then, many cases were reported from various states across the country. Pondicherry reported its first case in the month of March, 2020 and there reached a spike during August 2020 (Laxminarayan et al., 2020). Whole genome sequencing helps us in understanding the geographical frequency, adaptation of virus overtime, nature of disease transmission, pathogenesis, for vaccine and drug designing, vaccine dosage and hospitalization. Thus, we did a WGS from the UT of Puducherry including the representative samples (128) taken from all three waves. The UT is one of the tourist places and also French settlements are common. This results in the virus transmission from international travellers and also by local transmission like mass gatherings. The variants circulating in Puducherry during all three waves deciphered through WGS, was compared with the variants circulating globally and in India during the same time. In our study, we reported 14 Sub-lineages (B.1.560, B.1.1, B.1.1.354, B.1.351, B.1.1.7, B.1.617.2, B.1.617, AY.127, B.1.617.1, AY.59, BA.2, BA.2.17, BA.1.1, BA.1.1.7) belonging to lineage A and B.

They are the ancestral lineages of the later circulating lineages in different parts of the world. A and B are first sequenced lineage in January 2020, China (Tan et al., 2020). The lineages were similar to the one circulating in Europe, Australia, USA, UK, South Africa, Singapore, Malaysia, and Italy. The lineages in first wave belong to B.1.560, B.1.1, B.1.1.354 which are classified as uncommon variants according to WHO nomenclature. In second wave it was a combination of both A and B lineages with majority of them being B.1.617.2 (Delta variant). The interesting finding from our study is that many of the variants falls under uncommon category which does not fit either VOC or VOI as per WHO. Since these viruses were under circulation during the outbreak, what would be the role of these strains in terms of disease severity, pathogenesis and mutation needs detailed evaluations for effective disease surveillance and pandemic preparedness. In a multicentric study by ICMR (Gupta et al., 2021), totally 26 samples were collected from Puducherry region and the most common variant identified was Delta (B.1.617.2) followed by alpha and Kappa. They found two Delta variants AY.1 And AY.2 which had K417N spike protein mutation which was also responsible for immune escape and increase in infectivity. The most common variant over different regions of India was Delta while in northern region it was Alpha VOC. The breakthrough infection was common with Delta variant because of the reduced neutralizing capacity of the currently available vaccine and the high rate of transmission of Delta variant during that period (Gupta et al., 2021). A study from Gujarat was done with 502 sequenced samples from deceased and recovered patients and were compared nationally and globally. The missense mutation found in C28854T (Ser194Leu) among the deceased patients had an allele frequency of 47% and 7% respectively in Gujarat and Globally. It is a deleterious mutation in Nucleocapsid (N) gene. This shows a distinct mutation in Gujarat responsible for disease severity among the deceased (Joshi et al., 2021). This is followed by a report from Delhi, the capital state of India, in which 612 samples were sequenced spanning first three peaks of pandemic. The study reported 26 lineages with novel mutations detected across the 3 peaks (Gautam et al., 2022). Like ours this is also a single center study, in addition to their findings we have also included samples from Jan 2022 to Mar 2022, where Omicron type dominated with a total of 25. Similar to that study, we have also categorized our samples based on median CT. Though the minimum CT remained almost same in all the three peaks, the median CT was found to be less than 20 in peak three where Omicron is predominant type. In the first two peaks median CT values were found to be more than 20. This is in contrast to what has been reported by Gautam et al. 2022, where the median CT in the first two peaks were found to be 15 (Gautam et al., 2022).

The number of mutations per sample was found to be more during third peak followed by second peak. This could be due to units of mutation per site per year and the number of substitutions per site per replication cycle. Studies shows that the mutation rate as 10^-3^ mutations per site per year. This proves that the virus undergoes continuous replication which results in the accumulation of mutations over the course of the year (Urhan & Abeel, 2021; Vilar & Isom, 2021). The identified lineages in our study were found to vary with time. During the first peak B.1.560 lineage was found to be more common which was replaced by B.1.617.2 lineage during second peak. This was similar to a study in Delhi (Gautam et al., 2022). The beginning of third peak had B.1.617.2 lineage which slowly less detected with an increase in BA.2 and BA.1.1.7 lineage. This timely change in the mutation was reported across the world (Volz et al., 2021). The most common mutations observed was Spike_D614G. The mutation in spike protein has been well described by many studies (Guruprasad, 2021; Harvey et al., 2021; Magazine et al., 2022). ‘S’ protein of SARS-CoV-2 is responsible for target recognition, binding to ACE2 and entry into host cell. Frequent mutations in S proteins could have some impact on the binding properties to host ACE2 and RBD (Mariappan et al., 2020, 2022). For instance the D614G mutation observed in the current study was a kind of pervasive in the all the sequenced cases of COVID-19, later reported to alter the confirmation of RBD and enhances the affinity between ‘S’ and ACE2 (Gobeil et al., 2021; Magazine et al., 2022). Earlier studies have reported NSP2, 3 4 6 and 12 as recurrent hotspots in different geographic areas like Asia, Oceania, Europe and North America, whereas in the case of NSP13 helicase, mutants were reported only in North America (Pachetti et al., 2020). The new mutations detected in the present study includes NSP16_I171T, NS7a_I4J, NSP3_I1094S, NSP3_S925Y, NSP4_T370A, NSP5_A260D and Spike_N540T. Among them NSP16_I171T is detected among five patients (Alpha-2, Beta-1, kappa-1, delta-1) and remaining mutations in one patient each. These unique mutations were confirmed by fastq files generated by the Local Run Manager (Illumina) were aligned to the SARS-CoV-2 reference genome (NC_045512.2) using a Burrows-Wheeler Aligner (ver. 0.7.17) (Li & Durbin, 2009). The consensus sequence for each genome sequence was verified with NCoV-Tools. GATK pipeline was used for the detection of single nucleotide variants and short insertions/deletions with an average coverage of 1000x and minimum depth of 100x. SnpEff (ver. 5.0c) (Cingolani et al., 2012) was then used to annotate the filtered variants (VAF > 0.05). When the mutation frequencies were <0.05, whether the called reads truly existed was confirmed with an integrated genome viewer (IGV), CoVsurver and COVID-19 genome annotator.

The known mutations reported in the current study were mostly confined to NS1 to NS5 with NS3 topping the list. NS3 is considered an important non-structural protein (NSP) of the virus for its replication and regulating host protein synthesis. Among all the NSP’s (1- 16), NSP3 has been reported to have maximum number of mutations, however in vitro studies that validate the role of these mutations in affecting host protein synthesis or virus replication efficiency are still lacking. In the same note it would be intriguing to study extensively if the NS3 mutations observed in the current study deserves further studies. Another important NSP that are widely used by the virus for its replication and transcription is NSP16. NSP 16 encodes methyl transferase activities that are essential for virus mediated immune evasion. The S-Adenosylmethionine (SAM) dependent 2’-O-Methyltransferase enzyme with the conserved catalytic tetrad ([K46-D130-K170-E203]) from SARS-CoV-2 consists of 2 subunits: nsp16 (catalytic subunit) and nsp10 (stimulatory subunit) with nsp16 being activated by heterodimerization with nsp10. Thus, the enzymatic activity of NSP16 is attained with the support from NSP10. In other words, NSP16 is activated by NS10, otherwise remains nascent form. In the present study unique or new mutations in NSP 16 are observed in five samples. NSP16 which was not reported to have many mutations unlike NSP3 could be a potential therapeutic target for SARS-SoV2 control. However, the mutations identified in the current study putforth an important question on how the mutations in NS16 will have impact on NS10/NS16 complex? How the transferase activity of NSP16 will be affected and how it modifies the immune evasion mechanism of the virus remains unanswered. To further understand this, we did a computational analysis and found that the mutation NSP16_I171T observed in our study is very significant since it is within the proximity of the conserved catalytic tetrad. Interestingly, from the multiple sequence alignment of nsp16 protein sequences from different species, it can be observed that the point mutation NSP16_I171T has occurred within the pan-coronavirus highly conserved motif **(Fig. S7)**. The effect of the mutation on overall protein stability and dynamics was analyzed using DynaMut2 (Rodrigues et al., 2021) by introducing point mutation NSP16_I171T in the nsp16-nsp10-SAM complex (PDB ID: 6W61). A decrease in folding free energy (ΔΔ*G*) of -3.41 kcal/mol was observed. In other words, improved flexibility is observed in the mutated protein (nsp16_I171T) when compared to the native structure. Moreover, a non-polar amino acid, Isoleucine is substituted by Threonine a polar amino acid, with hydrogen bond forming potential. Given the proximity of the mutation site to the catalytic tetrad, improved flexibility NSP16_I171T mutation and hydrogen bond forming potential of Threonine, the mutation might probably have a strong influence in the catalytic activity of nsp16 (Rosas-Lemus et al., 2020).

We have considered deep sequencing technology to understand the genetic diversity of SARS-CoV-2 genome across the pandemic. We understand few limitations of the current study that include an unequal number of samples analysed during the peaks that prohibits us from understanding the distribution of variants based on age and sex. The significance of novel mutations identified, and its clinical correlations were not assessed. Also, since majority of the patients did not take single dose of vaccine at the time of sampling, the effect of mutants on vaccine immunology that are currently in usage are not addressed.

## Conclusion

The top mutations as observed in the current study led us to speculate that increased mutation rate in the 2^nd^ and 3^rd^ peaks might be due to the impaired proof-reading activity and altered polymerase activity of the virus. Thus, further studies are needed to (i) understand the molecular mechanisms that are involved in the increased mutational rate in key viral NSP’s (ii) check if the mutation in key NSP’s has resulted in increased viral replication and immune evasion (iii) study if the mutations resulted in resistance to current antivirals in use (iv) develop broad spectrum antivirals. Hence it is more important to study the evolutionary and genomic epidemiology of SARS-CoV-2 as it has a tendency to mutate very frequently. The continuous bio-surveillance involving detection of pathogens and new mutants using genome sequencing techniques and sharing of information at regional and global levels could limit the future outbreaks and pandemic.

## Data Availability

SARS-CoV-2 genome sequencing reads were deposited in the National Center for Biotechnology Information (NCBI - GenBank) with the accession number OP599771- OP599898.

## Ethics Approval

The study was approved by the Institute Research Committee (IRC) and Institute Human Ethics Committee (IHEC) of Mahatma Gandhi Medical College and Research Institute (MGMCRI/IRC/52/2021/04/IHEC/42).

## Consent to Participate

In this study de-identified and anonymized archived samples were subjected to whole-genome sequencing. No personal details except clinically relevant information (age, sex, symptoms, vaccination details, mode of quarantine details and outcome) were used for the purpose of the study. Hence written individual consent form was waived off by the Institutional Human Ethics Committee.

## Authors contribution

**VK:** Conceptualization, Supervision, Formal analysis, Writing – Original Draft, Writing – Review and Editing; **ABP:** Conceptualization, Supervision, Methodology, Writing – Original Draft, Writing – Review and Editing; **VM:** Formal analysis, Software, Writing – Review and Editing; **MR:** Formal analysis, Investigation; **PR**: Methodology, Resource, Writing –Review and Editing; **RR:** Data Curation, Formal analysis, Software; **BM:** Formal Analysis, Methodology, Software; **JME:** Resource, Writing – Review and Editing; **MV:** Data Curation, Formal analysis, Software, Writing – Review and Editing; **SRR:** Conceptualization, Supervision, funding acquisition

## Acknowledgment

The authors greatly acknowledge the funding support provided by “Sri Balaji Educational and Charitable Public Trust-SBECPT”, Sri Balaji Vidyapeeth to carry out the study. We acknowledge Dr. Pravin Charles, Dr. Namrata K Bhosale, Dr. Ramyapriyadarshini for their constant support during the study, Mrs. Humera Begum and Mrs.Tamizhmani for their technical support.

## Reference

Bartlow, A. W., Middlebrook, E. A., Romero, A. T., & Fair, J. M. (2021). How Cooperative Engagement Programs Strengthen Sequencing Capabilities for Biosurveillance and Outbreak Response. Frontiers in Public Health, 9, 648424. 10.3389/fpubh.2021.648424

Cingolani, P., Platts, A., Wang, L. L., Coon, M., Nguyen, T., Wang, L., Land, S. J., Lu, X., & Ruden, D. M. (2012). A program for annotating and predicting the effects of single nucleotide polymorphisms, SnpEff. Fly, 6(2), 80–92. 10.4161/fly.19695

Coumare, V. N., Pawar, S. J., Manoharan, P. S., Pajanivel, R., Shanmugam, L., Kumar, H., Boratne, A. V., Subramanian, B., Easow, J. M., Sivaprakash, B., Kalaivani, R., Renuka, K., Prabavathy, S., Angeline, K., Pillai, A. B., & Rao, S. R. (2021). COVID- 19 Pandemic-Frontline Experiences and Lessons Learned From a Tertiary Care Teaching Hospital at a Suburban Location of Southeastern India. Frontiers in Public Health, 9, 673536. 10.3389/fpubh.2021.673536

Cucinotta, D., & Vanelli, M. (2020). WHO Declares COVID-19 a Pandemic. Acta Bio Medica : Atenei Parmensis, 91(1), 157–160. 10.23750/abm.v91i1.9397

Faria, N. R., Mellan, T. A., Whittaker, C., Claro, I. M., Candido, D. da S., Mishra, S., Crispim, M. A. E., Sales, F. C. S., Hawryluk, I., McCrone, J. T., Hulswit, R. J. G., Franco, L. A. M., Ramundo, M. S., de Jesus, J. G., Andrade, P. S., Coletti, T. M., Ferreira, G. M., Silva, C. A. M., Manuli, E. R., … Sabino, E. C. (2021). Genomics and epidemiology of the P.1 SARS-CoV-2 lineage in Manaus, Brazil. Science (New York, N.Y.), 372(6544), 815–821. 10.1126/science.abh2644

Frampton, D., Rampling, T., Cross, A., Bailey, H., Heaney, J., Byott, M., Scott, R., Sconza, R., Price, J., Margaritis, M., Bergstrom, M., Spyer, M. J., Miralhes, P. B., Grant, P., Kirk, S., Valerio, C., Mangera, Z., Prabhahar, T., Moreno-Cuesta, J., … Nastouli, E. (2021). Genomic characteristics and clinical effect of the emergent SARS-CoV-2 B.1.1.7 lineage in London, UK: A whole-genome sequencing and hospital-based cohort study. The Lancet. Infectious Diseases, 21(9), 1246–1256. 10.1016/S1473-3099(21)00170-5

Gautam, P., Paul, D., Suroliya, V., Garg, R., Agarwal, R., Das, S., Kaur, U. S., Pandey, A., Bhugra, A., Tarai, B., Bihari, C., Sarin, S. K., & Gupta, E. (2022). SARS-CoV-2 Lineage Tracking, and Evolving Trends Seen during Three Consecutive Peaks of Infection in Delhi, India: A Clinico-Genomic Study. Microbiology Spectrum, 10(2), e02729-21. 10.1128/spectrum.02729-21

Gobeil, S. M.-C., Janowska, K., McDowell, S., Mansouri, K., Parks, R., Manne, K., Stalls, V., Kopp, M. F., Henderson, R., Edwards, R. J., Haynes, B. F., & Acharya, P. (2021). D614G Mutation Alters SARS-CoV-2 Spike Conformation and Enhances Protease Cleavage at the S1/S2 Junction. Cell Reports, 34(2), 108630. 10.1016/j.celrep.2020.108630

Gupta, N., Kaur, H., Yadav, P. D., Mukhopadhyay, L., Sahay, R. R., Kumar, A., Nyayanit, D. A., Shete, A. M., Patil, S., Majumdar, T., Rana, S., Gupta, S., Narayan, J., Vijay, N., Barde, P., Nataraj, G., B., A. K., Kumari, M. P., Biswas, D., … Abraham, P. (2021). Clinical Characterization and Genomic Analysis of Samples from COVID-19 Breakthrough Infections during the Second Wave among the Various States of India. Viruses, 13(9), 1782. 10.3390/v13091782

Guruprasad, L. (2021). Human SARS CoV-2 spike protein mutations. Proteins, 89(5), 569–576. 10.1002/prot.26042

Harvey, W. T., Carabelli, A. M., Jackson, B., Gupta, R. K., Thomson, E. C., Harrison, E. M., Ludden, C., Reeve, R., Rambaut, A., COVID-19 Genomics UK (COG-UK) Consortium, Peacock, S. J., & Robertson, D. L. (2021). SARS-CoV-2 variants, spike mutations and immune escape. Nature Reviews. Microbiology, 19(7), 409–424. 10.1038/s41579-021-00573-0

Joshi, M., Puvar, A., Kumar, D., Ansari, A., Pandya, M., Raval, J., Patel, Z., Trivedi, P., Gandhi, M., Pandya, L., Patel, K., Savaliya, N., Bagatharia, S., Kumar, S., & Joshi, C. (2021). Genomic Variations in SARS-CoV-2 Genomes From Gujarat: Underlying Role of Variants in Disease Epidemiology. Frontiers in Genetics, 12, 586569. 10.3389/fgene.2021.586569

Khandia, R., Singhal, S., Alqahtani, T., Kamal, M. A., El-Shall, N. A., Nainu, F., Desingu, P. A., & Dhama, K. (2022). Emergence of SARS-CoV-2 Omicron (B.1.1.529) variant, salient features, high global health concerns and strategies to counter it amid ongoing COVID-19 pandemic. Environmental Research, 209, 112816. 10.1016/j.envres.2022.112816

Laxminarayan, R., Wahl, B., Dudala, S. R., Gopal, K., Mohan B, C., Neelima, S., Jawahar Reddy, K. S., Radhakrishnan, J., & Lewnard, J. A. (2020). Epidemiology and transmission dynamics of COVID-19 in two Indian states. *Science (New York*, N.Y.), 370(6517), 691–697. 10.1126/science.abd7672

Letunic, I., & Bork, P. (2021). Interactive Tree Of Life (iTOL) v5: An online tool for phylogenetic tree display and annotation. Nucleic Acids Research, 49(W1), W293– W296. 10.1093/nar/gkab301

Li, H., & Durbin, R. (2009). Fast and accurate short read alignment with Burrows-Wheeler transform. *Bioinformatics (Oxford*, England), 25(14), 1754–1760. 10.1093/bioinformatics/btp324

Lyngse, F. P., Kirkeby, C. T., Denwood, M., Christiansen, L. E., Mølbak, K., Møller, C. H., Skov, R. L., Krause, T. G., Rasmussen, M., Sieber, R. N., Johannesen, T. B., Lillebaek, T., Fonager, J., Fomsgaard, A., Møller, F. T., Stegger, M., Overvad, M., Spiess, K., & Mortensen, L. H. (2022). Household transmission of SARS-CoV-2 Omicron variant of concern subvariants BA.1 and BA.2 in Denmark. Nature Communications, 13(1), 5760. 10.1038/s41467-022-33498-0

Magazine, N., Zhang, T., Wu, Y., McGee, M. C., Veggiani, G., & Huang, W. (2022). Mutations and Evolution of the SARS-CoV-2 Spike Protein. Viruses, 14(3), 640. 10.3390/v14030640

Mandal, S., Arinaminpathy, N., Bhargava, B., & Panda, S. (2021). Plausibility of a third wave of COVID-19 in India: A mathematical modelling based analysis. The Indian Journal of Medical Research, *153*(5 & 6), 522–532. 10.4103/ijmr.ijmr_1627_21

Mariappan, V., Ranganadin, P., Shanmugam, L., Rao, S. R., & Balakrishna Pillai, A. (2022). Early shedding of membrane-bounded ACE2 could be an indicator for disease severity in SARS-CoV-2. Biochimie, 201, 139–147. 10.1016/j.biochi.2022.06.005

Mariappan, V., S R, R., & Balakrishna Pillai, A. (2020). Angiotensin-converting enzyme 2: A protective factor in regulating disease virulence of SARS-COV-2. IUBMB Life, 72(12), 2533–2545. 10.1002/iub.2391

Minh, B. Q., Schmidt, H. A., Chernomor, O., Schrempf, D., Woodhams, M. D., von Haeseler, A., & Lanfear, R. (2020). IQ-TREE 2: New Models and Efficient Methods for Phylogenetic Inference in the Genomic Era. Molecular Biology and Evolution, 37(5), 1530–1534. 10.1093/molbev/msaa015

Pachetti, M., Marini, B., Benedetti, F., Giudici, F., Mauro, E., Storici, P., Masciovecchio, C., Angeletti, S., Ciccozzi, M., Gallo, R. C., Zella, D., & Ippodrino, R. (2020). Emerging SARS-CoV-2 mutation hot spots include a novel RNA-dependent-RNA polymerase variant. Journal of Translational Medicine, 18(1), 179. 10.1186/s12967-020-02344-6

Rodrigues, C. H. M., Pires, D. E. V., & Ascher, D. B. (2021). DynaMut2: Assessing changes in stability and flexibility upon single and multiple point missense mutations. Protein Science: A Publication of the Protein Society, 30(1), 60–69. 10.1002/pro.3942

Rosas-Lemus, M., Minasov, G., Shuvalova, L., Inniss, N. L., Kiryukhina, O., Wiersum, G., Kim, Y., Jedrzejczak, R., Maltseva, N. I., Endres, M., Jaroszewski, L., Godzik, A., Joachimiak, A., & Satchell, K. J. F. (2020). The crystal structure of nsp10-nsp16 heterodimer from SARS-CoV-2 in complex with S-adenosylmethionine. bioRxiv, 2020.04.17.047498. 10.1101/2020.04.17.047498

Sahebi, S., & Keikha, M. (2022). Clinical features of SARS-CoV-2 Omicron BA.2; Lessons from previous observations – Correspondence. International Journal of Surgery (London, England), 104, 106754. 10.1016/j.ijsu.2022.106754

Sarkar, R., Mitra, S., Chandra, P., Saha, P., Banerjee, A., Dutta, S., & Chawla-Sarkar, M. (2021). Comprehensive analysis of genomic diversity of SARS-CoV-2 in different geographic regions of India: An endeavour to classify Indian SARS-CoV-2 strains on the basis of co-existing mutations. Archives of Virology, 166(3), 801–812. 10.1007/s00705-020-04911-0

Tan, W., Zhao, X., Ma, X., Wang, W., Niu, P., Xu, W., F. Gao, G., Wu, G., MHC Key Laboratory of Biosafety, National Institute for Viral Disease Control and Prevention, China CDC, Beijing, China, & Center for Biosafety Mega-Science, Chinese Academy of Sciences, Beijing, China. (2020). A Novel Coronavirus Genome Identified in a Cluster of Pneumonia Cases—Wuhan, China 2019−2020. China CDC Weekly, 2(4), 61–62. 10.46234/ccdcw2020.017

Tegally, H., Wilkinson, E., Giovanetti, M., Iranzadeh, A., Fonseca, V., Giandhari, J., Doolabh, D., Pillay, S., San, E. J., Msomi, N., Mlisana, K., von Gottberg, A., Walaza, S., Allam, M., Ismail, A., Mohale, T., Glass, A. J., Engelbrecht, S., Van Zyl, G., … de Oliveira, T. (2021). Detection of a SARS-CoV-2 variant of concern in South Africa. Nature, 592(7854), 438–443. 10.1038/s41586-021-03402-9

Thakur, V., Bhola, S., Thakur, P., Patel, S. K. S., Kulshrestha, S., Ratho, R. K., & Kumar, P. (2022). Waves and variants of SARS-CoV-2: Understanding the causes and effect of the COVID-19 catastrophe. Infection, 50(2), 309–325. 10.1007/s15010-021-01734-2

Urhan, A., & Abeel, T. (2021). Emergence of novel SARS-CoV-2 variants in the Netherlands. Scientific Reports, 11(1), 6625. 10.1038/s41598-021-85363-7

Vilar, S., & Isom, D. G. (2021). One Year of SARS-CoV-2: How Much Has the Virus Changed? Biology, 10(2), 91. 10.3390/biology10020091

Volz, E., Mishra, S., Chand, M., Barrett, J. C., Johnson, R., Geidelberg, L., Hinsley, W. R., Laydon, D. J., Dabrera, G., O’Toole, Á., Amato, R., Ragonnet-Cronin, M., Harrison, I., Jackson, B., Ariani, C. V., Boyd, O., Loman, N. J., McCrone, J. T., Gonçalves, S., … Ferguson, N. M. (2021). Assessing transmissibility of SARS-CoV-2 lineage B.1.1.7 in England. Nature, 593(7858), 266–269. 10.1038/s41586-021-03470-x

Yadav, P. D., Potdar, V. A., Choudhary, M. L., Nyayanit, D. A., Agrawal, M., Jadhav, S. M., Majumdar, T. D., Shete-Aich, A., Basu, A., Abraham, P., & Cherian, S. S. (2020). Full-genome sequences of the first two SARS-CoV-2 viruses from India. The Indian Journal of Medical Research, *151*(2 & 3), 200–209. 10.4103/ijmr.IJMR_663_20

