## Supplement tables and figures for "GENOMIC PROFILING OF SARS-COV-2 STRAINS CIRCULATING IN SOUTH EASTERN REGION OF INDIA DURING THREE WAVES OF PANDEMIC"

### List of Supplementary Tables

**Table S1. Variants from peak to peak**

| Variants | 2020 | | 2021 | | | | 2022 |
|---|---|---|---|---|---|---|---|
| | Aug-Sep | Oct-Nov | Jan-Mar | Apr-Jun | Jul-Sep | Oct-Nov | Jan-Mar |
| **VOC_Alpha** | 0 | 0 | 2 | 5 | 0 | 0 | 0 |
| **VOC_Beta** | 0 | 0 | 1 | 3 | 0 | 0 | 0 |
| **VOI_Kappa** | 0 | 0 | 0 | 4 | 0 | 0 | 0 |
| **VOC_Delta** | 0 | 0 | 0 | 55 | 8 | 4 | 0 |
| **VOC_Omicron** | 0 | 0 | 0 | 0 | 0 | 0 | 25 |
| **Others** | 10 | 7 | 0 | 3 | 1 | 0 | 0 |

<u>**List of Supplementary Figures**</u>

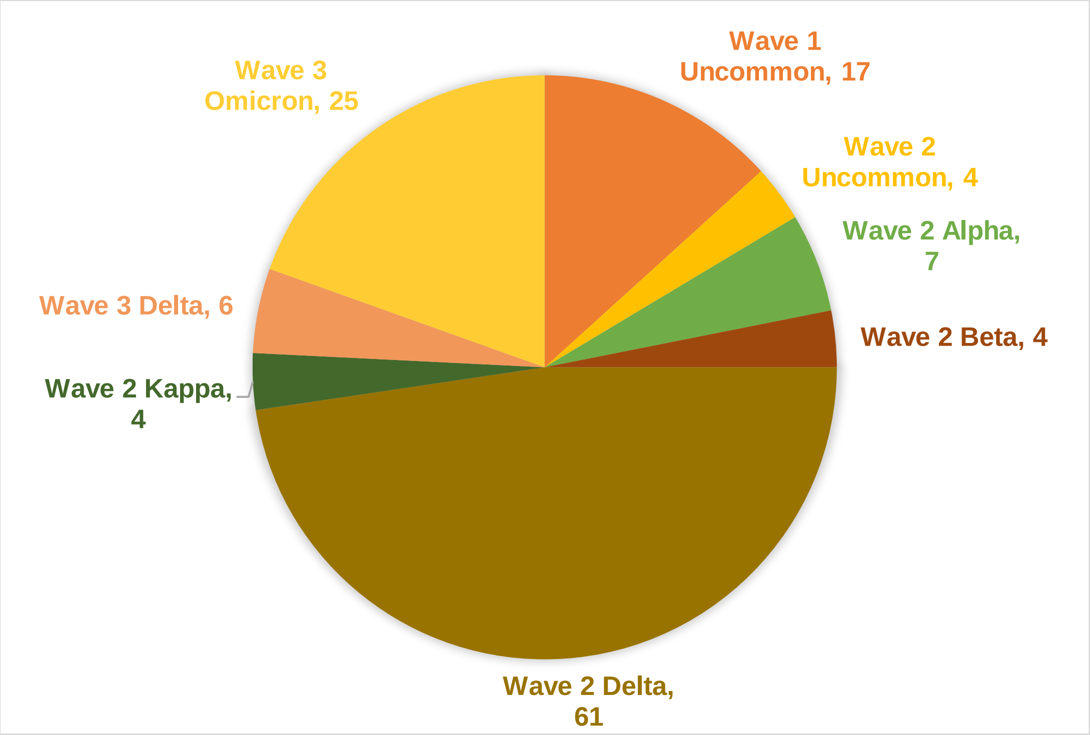


**Fig. S1. Distribution of Variants during three consecutive peaks**

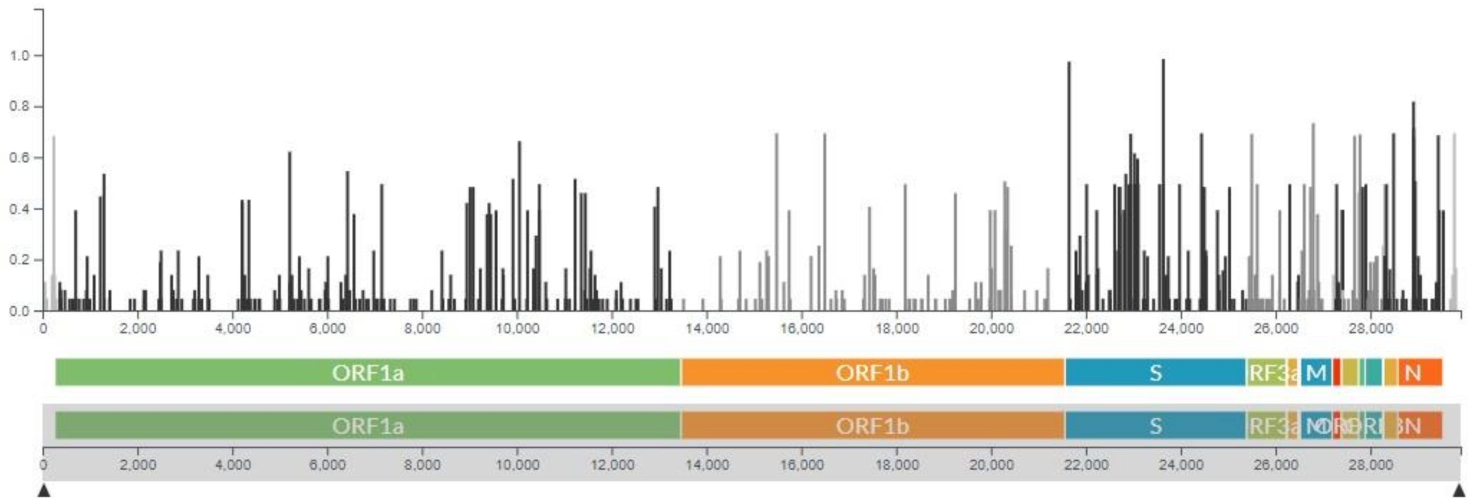


**Fig S2. COVID-19 Genome wide coverage**

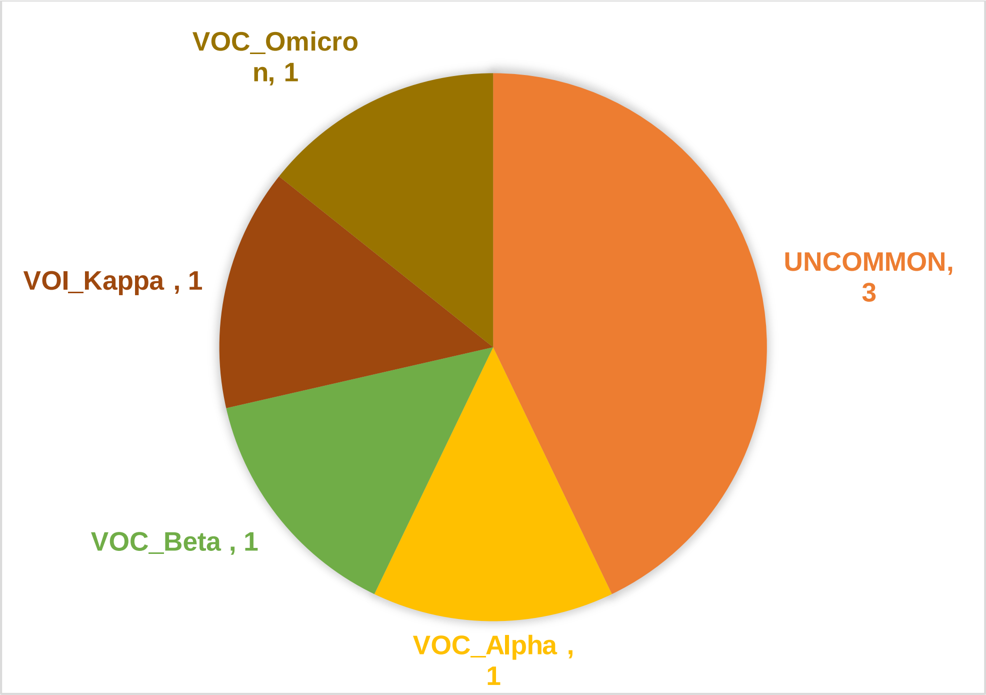


**Fig S3. Breakthrough Infection in Second and Third wave**

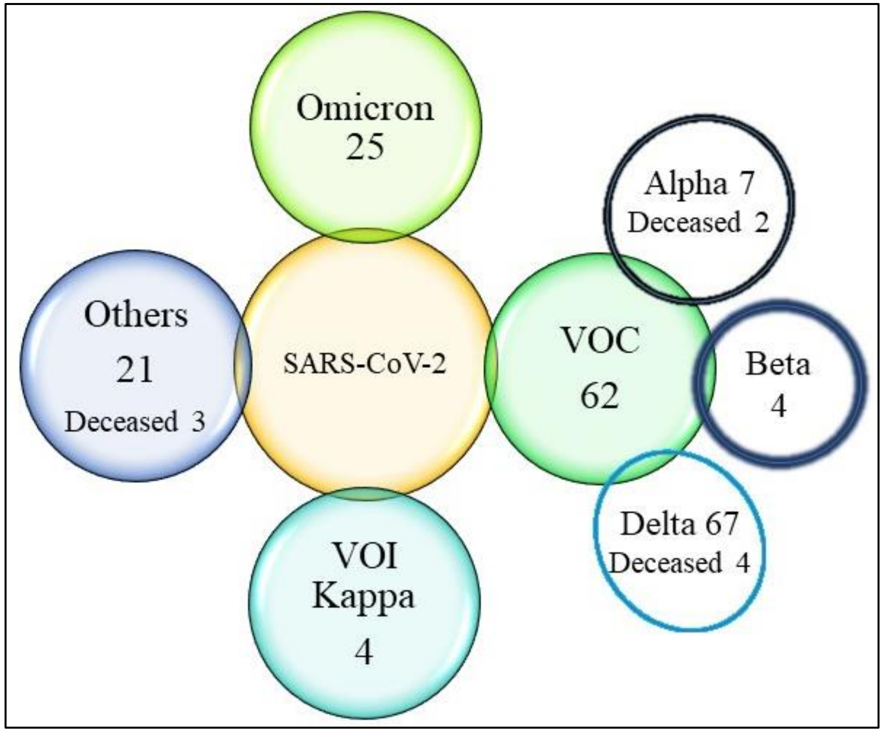


**Fig S4. Overview of total variants and the outcome**

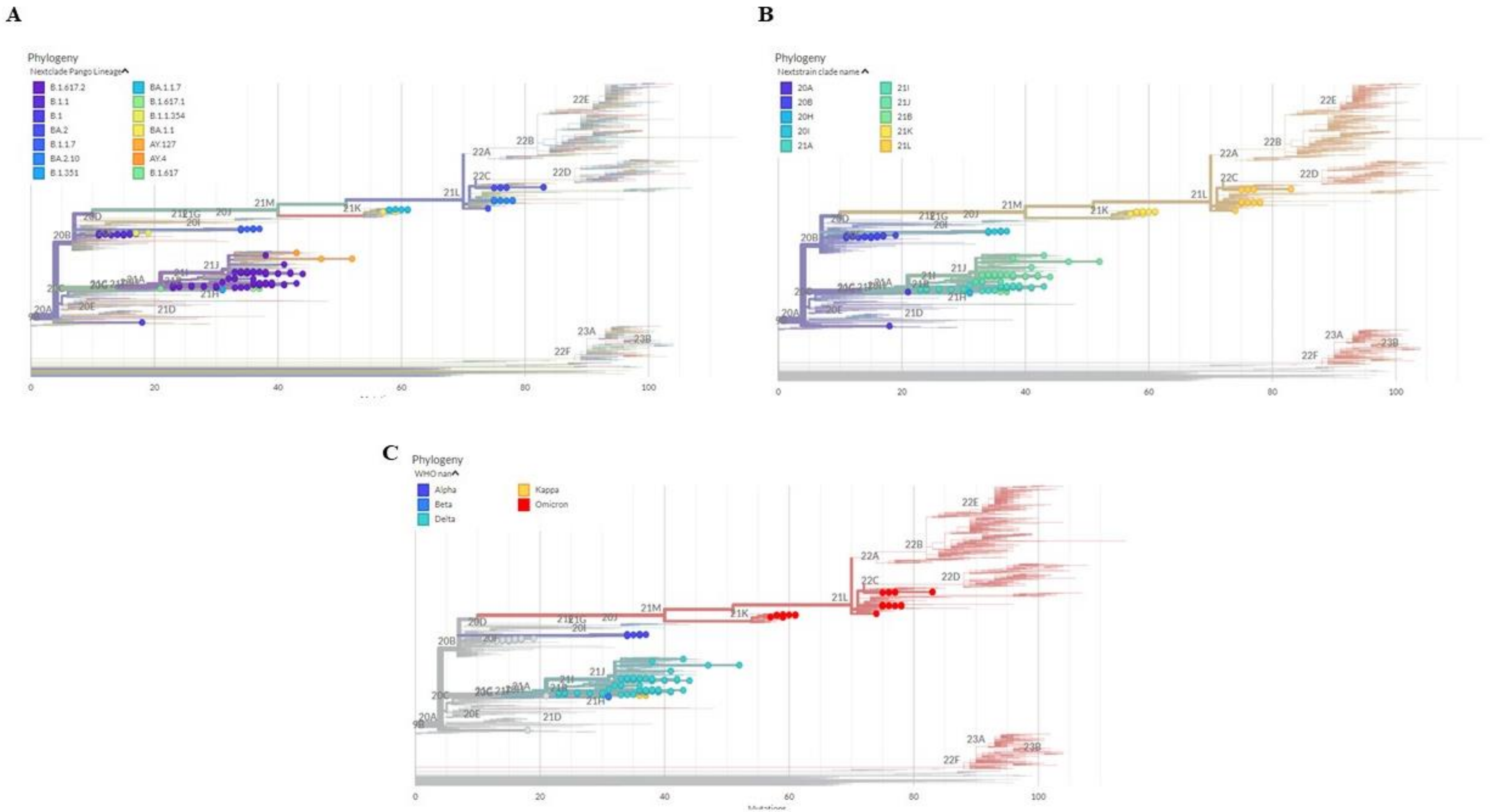


**Fig S5. Phylogenetic analysis of 128 SARS-CoV-2 patient samples in Puducherry from August 2020 to March 2022.** Dendrogram revealing the phylogenetic relationship between the SARS-CoV-2 genomes from 128 patient samples in Puducherry, aligned against the Wuhan Hu 1 (NC_045512.2) reference genome. The dendrogram was generated using FastTree, which infers approximately-maximum-likelihood phylogenetic trees from alignments of nucleotides. Major clades from pangolin lineages, Nextstrain and WHO lineages are indicated at the branch points.

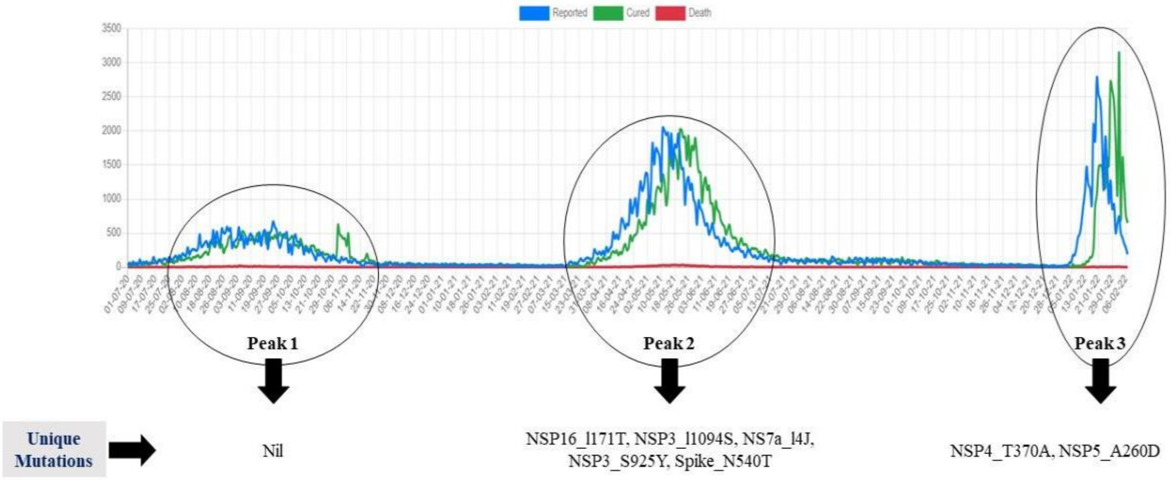


**Fig S6. Distribution of unique mutants during three consecutive peaks in Puducherry.** The sample collection during the peak 1 and peak 2 is similar to corresponding circle in the given image. Whereas the sample collection peak 3 was done during Sep 2021-Feb 2022.

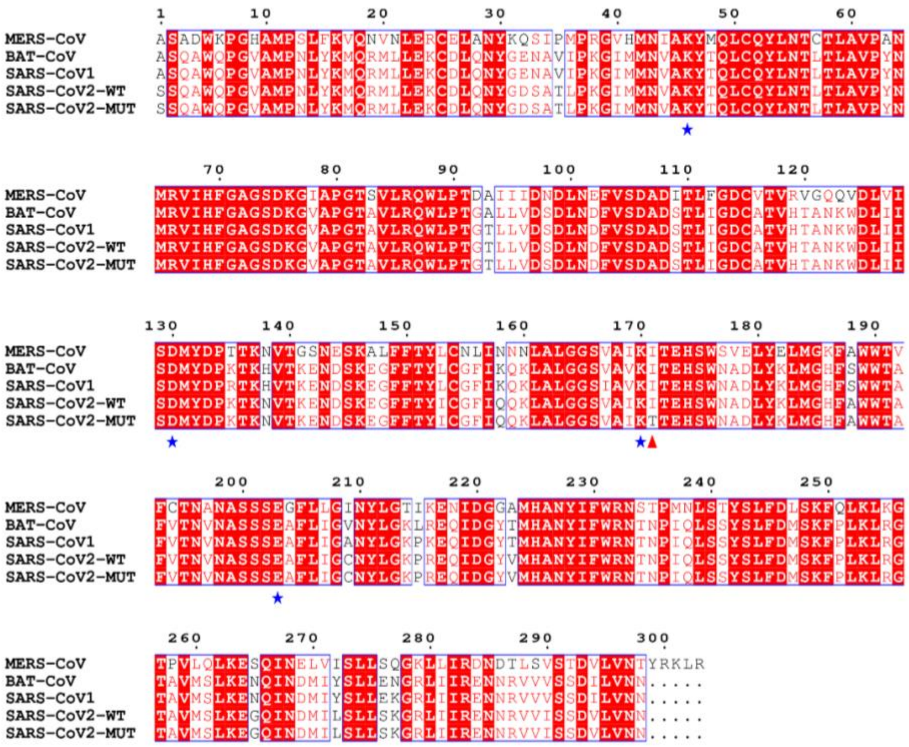


**Fig. S7**: Multiple Sequence Alignment of nsp16 protein sequences from different sources (MERS-CoV: UniProt ID K9N7C7, Bat-CoV: UniProt ID P0C6V9, SARS-CoV1: UniProt ID P0C6X7, SARS-CoV2: UniProt ID P0DTD1) as indicated in the sequence names. The catalytic tetrad residues are indicated as blue stars and the mutation site is indicated as red triangle.
